# Determinants of cerebral blood flow and arterial transit time in healthy older adults

**DOI:** 10.1101/2023.12.13.571578

**Authors:** Jack Feron, Katrien Segaert, Foyzul Rahman, Sindre H Fosstveit, Kelsey E Joyce, Ahmed Gilani, Hilde Lohne-Seiler, Sveinung Berntsen, Karen J Mullinger, Samuel J E Lucas

## Abstract

Cerebral blood flow (CBF) and arterial transit time (ATT), markers of brain vascular health, worsen with age. The primary aim of this cross-sectional study was to identify modifiable determinants of CBF and ATT in healthy older adults (n=78, aged 60–81 yrs). Associations between cardiorespiratory fitness and CBF or ATT were of particular interest as the impact of cardiorespiratory fitness is not clear in existing literature. Secondly, this study assessed whether CBF or ATT relate to cognitive function in older adults. Results from multiple linear regressions found higher BMI was associated with lower global CBF (β=-0.35, *P*=0.008) and a longer global ATT (β=0.30, P=0.017), global ATT lengthened with increasing age (β=0.43, *P*=0.004), and higher cardiorespiratory fitness was associated with longer ATT in parietal (β=0.44, *P*=0.004) and occipital (β=0.45, *P*=0.003) regions. Global or regional CBF or ATT were not associated with processing speed, working memory, or attention. In conclusion, preventing excessive weight gain may help attenuate age-related declines in brain vascular health. ATT may be more sensitive to age-related decline than CBF, and therefore useful for early detection and management of cerebrovascular impairment. Finally, cardiorespiratory fitness appears to have little effect on CBF but may induce longer ATT in specific regions.

## Introduction

The global population aged over 60 is expected to almost double by 2050.^1^ Maintaining brain health is essential to healthy ageing. Cerebral blood flow (CBF) and arterial transit time (ATT) are markers of brain vascular health^2–4^ yet their determining factors are poorly understood in older adults, partly due to variations in measurement techniques, with few arterial spin labelling (ASL) studies accounting for differences in ATT when calculating CBF. The present study aimed to identify modifiable determinants of CBF and ATT in healthy older adults, accurately measured using ASL with multiple post-label delays, and establish whether CBF and ATT are associated with cognitive function.

Cognitive function is positively associated with independent living and quality of life in older adults,^5,6^ but progressively, and unavoidably, declines with age.^7^ Blood flow to and within the brain is important for brain health since the brain has a high relative energy consumption but lacks intracellular energy stores, meaning hypoperfusion can result in cell death and neuronal dysfunction.^8,9^ Higher CBF has been linked with superior cognitive function in healthy older adults,^10–14^ although not all studies converge.^12,15–17^ It is well established that CBF declines with age^4,18,19^ and previous research in older adults has already identified some modifiable determinants of CBF. For example, CBF is reported to be greater in older adults who are physically active,^20–22^ engage with social or leisure activities,^23^ have a lower BMI,^24–26^ have lower blood pressure,^19,26^ or have more elastic arteries.^27,28^ CBF in older adults thus appears to be impacted by physical activity, social interaction, body weight/composition, and cardiovascular health, which are all modifiable risk factors that can be addressed with simple lifestyle changes.

In the present study, cardiorespiratory fitness was of particular interest as a modifiable lifestyle factor that may impact CBF in older adults. This is because despite cardiorespiratory fitness’s well-documented benefits for cognition,^29–31^ its impact on CBF is unclear. Note that being physically active and having a high cardiorespiratory fitness are often assumed to coincide, however, cardiorespiratory fitness has a large genetic component,^32^ which may explain some of the inconsistencies in the literature around cardiorespiratory fitness and its relation to CBF. Nevertheless, maintaining higher cardiorespiratory fitness or physical activity levels is suggested to attenuate age-related decline in CBF^21,33^ and blood velocity within a cerebral artery (CBv),^34^ considered a proxy for CBF.^35^ Some cross-sectional research in older adults reports a positive association between cardiorespiratory fitness and CBF^36–39^ or CBv.^40^ Furthermore, aerobic exercise training in older adults has been shown to increase CBF^28,41–45^ or CBv.^46–49^ In contrast, a negative association between cardiorespiratory fitness and CBF has also been found^50,51^ or a complete lack of association between cardiorespiratory fitness and CBF^52–54^ or CBv.^55–59^ Interestingly, reported changes in CBF are not usually global, but instead confined to specific regions of the brain.^33,43–45,50^ Moreover, regions are often small or only a portion of regions investigated show these associations.^38,43,44,51^ Of note, CBF and CBv are assessed from distinct regions of the cerebral vascular tree, which may partly explain discrepancies in results, alongside differences in the arteries or grey matter regions investigated, and variations in ASL protocols used for CBF measures. Furthermore, none of the aforementioned ASL studies account for ATT differences when calculating CBF. Given the complexity and inconsistency in the literature to date, more work is needed to understand the relationship between these variables, and with improved methodological approaches to better consider global versus regional CBF changes.

ATT is the time taken for blood to travel from large arteries in the neck to the cerebral tissue. ATT is primarily used to improve the quantification of CBF by adjusting for individual and regional differences in ATT, this is particularly important for older adults who experience longer ATT.^60^ However, measuring at multiple post-label delays with ASL to allow ATT as well as CBF to be calculated involves a longer MRI data acquisition and therefore relatively few studies investigate ATT as well as CBF. This means accuracy of CBF measurements are compromised in most ageing studies using ASL, where changes in ATT are expected, and that the determinants of ATT remain poorly understood. Like CBF, ATT is considered a marker of brain health whereby a shorter ATT indicates superior brain health. Indeed, research has shown that ATT lengthens with age,^4,61,62^ cerebral artery stenosis,^63^ and Alzheimer’s disease.^3^ Furthermore, prolonged ATT has been associated with cognitive impairment.^3,64^ Regarding cardiorespiratory fitness, a small study in older adults found no relationship with ATT (n=14), but a positive relationship in younger adults (n=18).^52^ Moreover, a lower BMI has been associated with a longer regional ATT in male older adults with coronary artery disease.^65^ These latter findings somewhat contradict the idea that a shorter ATT is indicative of better brain health. Further research is warranted to identify factors that affect ATT and to establish what is a ‘healthy’ ATT.

Therefore, the purpose of this cross-sectional study was to improve understanding of the determinants of resting grey matter CBF and ATT, both globally and regionally, in healthy older adults, with a particular focus on cardiorespiratory fitness, and whether CBF or ATT are associated with cognitive function. CBF and ATT were measured using ASL with multiple post-label delays, allowing adjustment of CBF for ATT differences. It was hypothesised that markers of superior general health (i.e., higher cardiorespiratory fitness/handgrip strength/grey matter volume or lower age/BMI/blood pressure) and cognitive function would be associated with greater CBF and a shorter ATT.

## Material and methods

### Study design

The data for this publication were collected as part of a larger study, The FAB Project (preregistration: https://osf.io/6fqg7, materials and data: https://osf.io/d7aw2/). The study was approved by the STEM Ethical Review Committee at the University of Birmingham (ERN_20-1107).

Participants were screened for eligibility before completing three experimental sessions on different days. Figure 1 shows the key outcome measures and desired session order (achieved for 87% of participants, with all experimental sessions completed within 5.2±3.2 weeks, and with 13±15 days between MRI and exercise sessions (<30 days for 91%).

**Figure 1:**
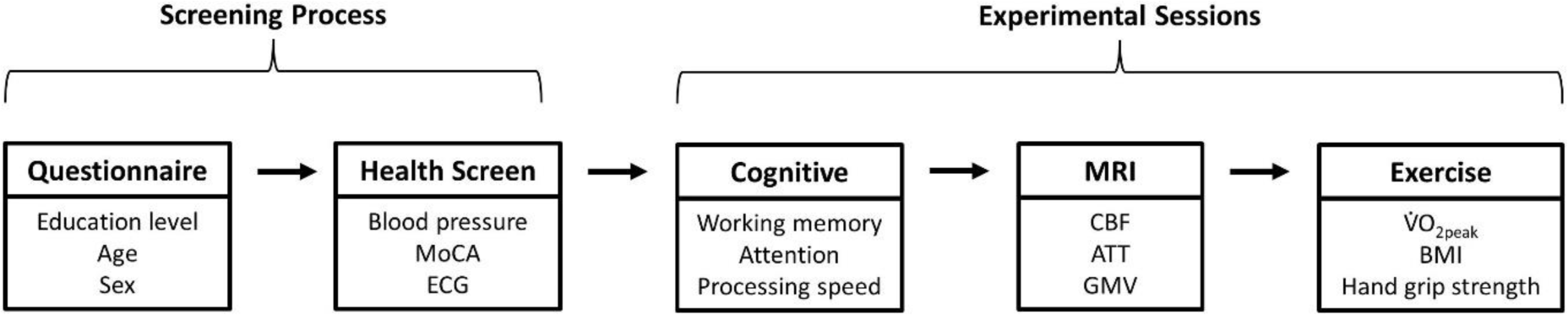
Flow chart of the screening and experimental sessions. MoCA; Montreal Cognitive Assessment, ECG; electrocardiogram MRI; magnetic resonance imaging, CBF; cerebral blood flow, ATT, arterial transit time, GMV; grey matter volume, *VO*_2*peak*_; peak oxygen consumption, BMI; body mass index.

### Participants

Ninety-four healthy older adults (aged 60 – 81 yrs) were enrolled. Participants were normotensive, cognitively normal, without historic or current diagnosis of serious health conditions, non-smokers, and self-reported to not meet recommended global activity guidelines.^66^ Section 1 of the supplementary material contains detailed inclusion criteria. MRI data was missing or unusable for n=16, leaving n=78 for analyses presented in this study. Participant characteristics are shown in Table 1.

**Table 1:** Participant characteristics.

|  | Total | Male | Female |
| --- | --- | --- | --- |
| n | 78 | 39 | 39 |
| Age (years) | 65±5 | 65±5 | 66±5 |
| Education (%) |  |  |  |
| Compulsory | 28 | 31 | 26 |
| Further | 33 | 33 | 33 |
| Undergraduate | 19 | 18 | 21 |
| Postgraduate | 19 | 18 | 21 |
| SBP (mmHg) | 140±13 | 140±12 | 139±15 |
| DBP (mmHg) | 82±8 | 83±7 | 82±8 |
| BMI (kg/m<sup>2</sup>) | 27±4 | 28±3 | 26±4 |
| $\dot{V}O_{2peak}$ (mL/kg/min) | 28±4 | 30±4 | 25±3 |

Values represent means ± standard deviation. SBP; systolic blood pressure, DBP; diastolic blood pressure, BMI; body mass index, $\dot{V}O_{2peak}$ ; peak oxygen consumption.

### Health screening: electrocardiogram (ECG), blood pressure, and cognitive impairment

Participants completed a resting 12-lead ECG (Cardiosoft, Vyaire, USA), three resting blood measurements (705IT, Omron, Japan), and the Montreal Cognitive Assessment (MoCA). Participants were excluded for severe ECG abnormalities, MoCA scores <23,^67^ and systolic/diastolic blood pressure of >160/>90 mmHg. Excluded participants were referred to their GP.

### Outcome Measures

#### Cardiorespiratory fitness

Participants completed an incremental exercise test on a treadmill (Pulsar 3p, H/P/Cosmos, Germany). Respiratory gases (*VO*_2_; oxygen consumption, *VCO*_2_; carbon dioxide production) were recorded continuously using a facemask (7450 V2, Hans Rudolph, USA) and metabolic cart (JAEGER Vyntus CPX, Vyaire, USA), as was heart rate and rhythm using a 12-lead ECG (Cardiosoft, Vyaire, USA). Rating of perceived exertion (RPE)^68^ and finger-prick blood [lactate] (Biosen C-Line, EKF Diagnostics, United Kingdom) were measured between stages. Stages were 4 min with a 1 min rest period between each stage. Treadmill speed started and remained at 3.8 km/h until either all possible elevation stages were completed (4, 7, 10, 13, 16, 19, and 20% gradient) or individual lactate threshold was reached (2.1 mmol/L increase over the mean of the two lowest values^69^). If all elevation stages were completed, 4 min stages continued with speed increasing 0.5 km/h per stage until lactate threshold. After reaching lactate threshold, 1 min stages were completed where speed increased 0.5 km/h per stage (rest periods were removed). Figure S1 shows a treadmill test format example.

Participants were asked to exercise to volitional exhaustion unless halted by the researcher due to ECG abnormalities or injury. Cardiorespiratory fitness was determined using peak oxygen consumption (*VO*_2*peak*_) (i.e., mean of the two highest 30-s intervals). Nine participants completed a sub-maximal test, *VO*_2*peak*_ was thus predicted using individual sub-maximal *VO*_2_ and heart rate data acquired from three of the first possible six stages using a linear regression. Full details and example (Figure S2) of the prediction method can be found in Section 2 of the supplementary material.

#### Hand grip strength

Hand grip dynamometer (5001, Takei, Japan) was adjusted for grip size in their dominant hand. Participants stood upright, arms by their sides, and squeezed the dynamometer as hard as possible, maintaining elbow extension and limiting shoulder movement. The highest score from three attempts was taken.

#### Cognitive function

*Working memory*: In a 2-back task, participants were presented a 3x3 grid. The stimulus was a single white square that continuously appears, disappears, and then reappears in one of the grid squares at random (n=60 trials, 1 s each). Participants identified when the white square appeared in the same location as it did two trials prior. Trials were excluded from analysis if incorrect or if response time <200 ms or greater than two standard deviations above/below the mean per participant. The primary outcome measure was *d* prime (*d*’), a measure of discriminability, a greater *d*’ indicates superior performance.

#### Attentional Network Task (ANT)

The computerised ANT assessed orienting, alerting, and executive control. The stimulus is a row of five arrows, each pointing left or right. As fast and as accurately as possible, participants reported the direction of the centre arrow using the left and right arrow keys. A central fixation cross is displayed for 400 ms, then a fixation cross (500 ms) and cue (100 ms) are presented simultaneously, and then only the fixation cross is displayed for a further 400 ms. A stimulus is then shown for a maximum of 1700 ms. The centre arrow can be congruent or incongruent (i.e., pointing in the same or opposite direction as the flankers, respectively; n=96 each), or neutral (i.e., central arrow flanked by target-irrelevant black blocks, n=96). The stimulus can appear above or below the fixation cross, cued by a black square (n=216) or not cued (n=72). There are three cue conditions: a spatial cue, a centre cue, or a double cue (n=72 each). The spatial cue indicates if the stimulus appears above or below the fixation cross, whereas the stimulus location remains ambiguous for the centre and the double cue. Twelve practice trials are followed by three blocks of 96 trials.

Trials were excluded from analysis if incorrect or if response time <200 ms or greater than two standard deviations above/below the mean per participant. Alerting scores were calculated as the no cue minus the double cue; Orienting scores by centre cue minus the spatial cue; Executive control scores were calculated by the incongruent target minus the congruent target (all for correct responses). High condition difference scores for alerting and orienting, and low condition difference scores for executive control, indicate better performance.

#### Processing speed

In a letter comparison task, participants were simultaneously presented with two strings of letters at the top and bottom of the screen, for a maximum of 2500 ms after presentation of a fixation cross (1000 ms). Strings were three or six characters long (n=48 trials, 24 each). As fast and as accurately as possible, participants identified whether the strings were the same or different. Mean response time and accuracy were calculated for each participant using data from trials involving only six-character strings. Trials were excluded from analysis if incorrect or if response time <200 ms or greater than two standard deviations above/below the mean per participant.

#### MRI data acquisition and analysis

An MRI scan session included structural, functional, and arterial spin labelling (ASL) scans. We used a 3-T system (MAGNETOM Prisma, Siemens, Germany) with 32-channel receiver head coil. Here, the focus is the ASL data and related scans, analysis of other data acquired can be found elsewhere.^70^ CBF and ATT data were collected using pseudo-continuous ASL scan with 3D GRASE readout (17:22 mins),^71,72^ see also Acknowledgements. ASL measurements are reproducible and agree well with values obtained using [^15^O]-water positron emission tomography (PET),^73^ considered the ‘gold-standard’ technique.

ASL imaging parameters were: repetition time (TR)=4100 ms, echo time (TE)=30.56 ms, in-plane resolution=3.5 mm^2^, slice thickness=3.5 mm, transversal slices=32, field of view (FOV)=224x224 mm, labelling duration=1508 ms, and post-label delays (PLDs)=200, 975, 1425, 1850, 2025, 2150, 2250, and 2300 ms. Four and twelve volumes of data were acquired for PLDs of 200–2250 ms and 2300 ms, respectively. PLD times and number of volumes acquired were optimised according to recomendations.^74^ Slices were positioned axially from the motor cortex and angled anterior-posterior in line with the participant’s anterior-posterior commissure (ACPC). A calibration M0 scan was acquired using these same parameters with the PLD set to 2000 ms. The T1-weighted structural scan (4:54 mins) was acquired to facilitate data analysis including, normalisation to a standard template brain and differentiation of grey and white matter. Structural T1-weighted (MPRAGE) imaging parameters were: TE=2.03 ms, TR=2000 ms, voxel size=1 mm^3^, sagittal slices=208, FOV=256 mm, and flip angle=8°.

ASL data were processed using the Oxford ASL toolbox (https://oxasl.readthedocs.io/en/latest/), which uses the FSL FABBER ASL package and Bayesian Inference to invert the kinetic model for ASL MRI (BASIL) to compute CBF and ATT maps.^75–77^ Parameters input to the kinetic models to estimate CBF and ATT were: bolus duration=1.508 s, tissue T1=1.3 s, arterial blood T1=1.65 s, labelling efficiency=0.85, inversion times=1708, 24838, 29338, 33588, 35338, 36588, 37588, and 38088 ms, all other input parameters were kept with default settings appropriate to pCASL acquisition. Partial volume error correction and adaptive spatial smoothing of the perfusion maps was performed using default settings in oxford_asl.^76,78^

Global and regional analysis was performed, assessed in native (individual participant) and MNI space, respectively. All CBF and ATT values refer to grey matter only. Regions of interest (ROI) were the cingulate gyrus and frontal, parietal, temporal, occipital, and motor cortices (Figure S3). The chosen ROIs have been used previously,^52^ and were broad because there were no specific *a priori* hypotheses of regions that would be affected by determinants or associated with cognitive function. MNI registration was poor for n=1, leaving n=77 for regional analysis. Additional information regarding data quality assessment and grey matter mask configuration can be found in Section 3 of the supplementary material.

Grey matter volume was estimated from structural T1 anatomical scans. Brain extraction tool (BET) removed non-brain tissue^79^ before segmentation of tissue types using FMRIB’s Automated Segmentation Tool (FAST).^80^

### Statistical Analysis

All statistics used multiple linear regressions (SPSS Statistics v.29, IBM, USA). To identify determinants, global CBF or ATT were the dependent variable with age, sex, blood pressure (systolic and diastolic), BMI, *VO*_2*peak*_, hand grip strength, and grey mater volume as independent variables. To assess regional associations with cardiorespiratory fitness, mean CBF or ATT of each of the six ROIs were the dependent variable with age, sex, *VO*_2*peak*_, and any other significant determinants of global CBF or ATT identified from the above analysis as independent variables. To identify associations with cognitive function, global CBF or ATT were the dependent variable with age, sex, education, and scores for processing speed (accuracy and response time), working memory (*d’* prime), and the three attentional domains (alerting, orienting, and executive control scores) as independent variables.

Cognitive data was missing for n=2, leaving n=76 for global analysis. The same analyses were performed using regional data, regional analysis details and results are shown in Section 5 of the supplementary material.

## Results

Mean global and regional values for CBF and ATT can be found in Table S1 and Figure S4.

### Determinants of global CBF

The overall global CBF (gCBF) model was not significant (R^2^ =0.38, F(8,69)=1.38, *P*=0.22). Of the eight independent variables, BMI was the only significant determinant of gCBF (β=-0.35, *P*=0.008; Figure 2), whereby gCBF decreased with increasing BMI. Data shown in Table 2.

**Figure 2:**
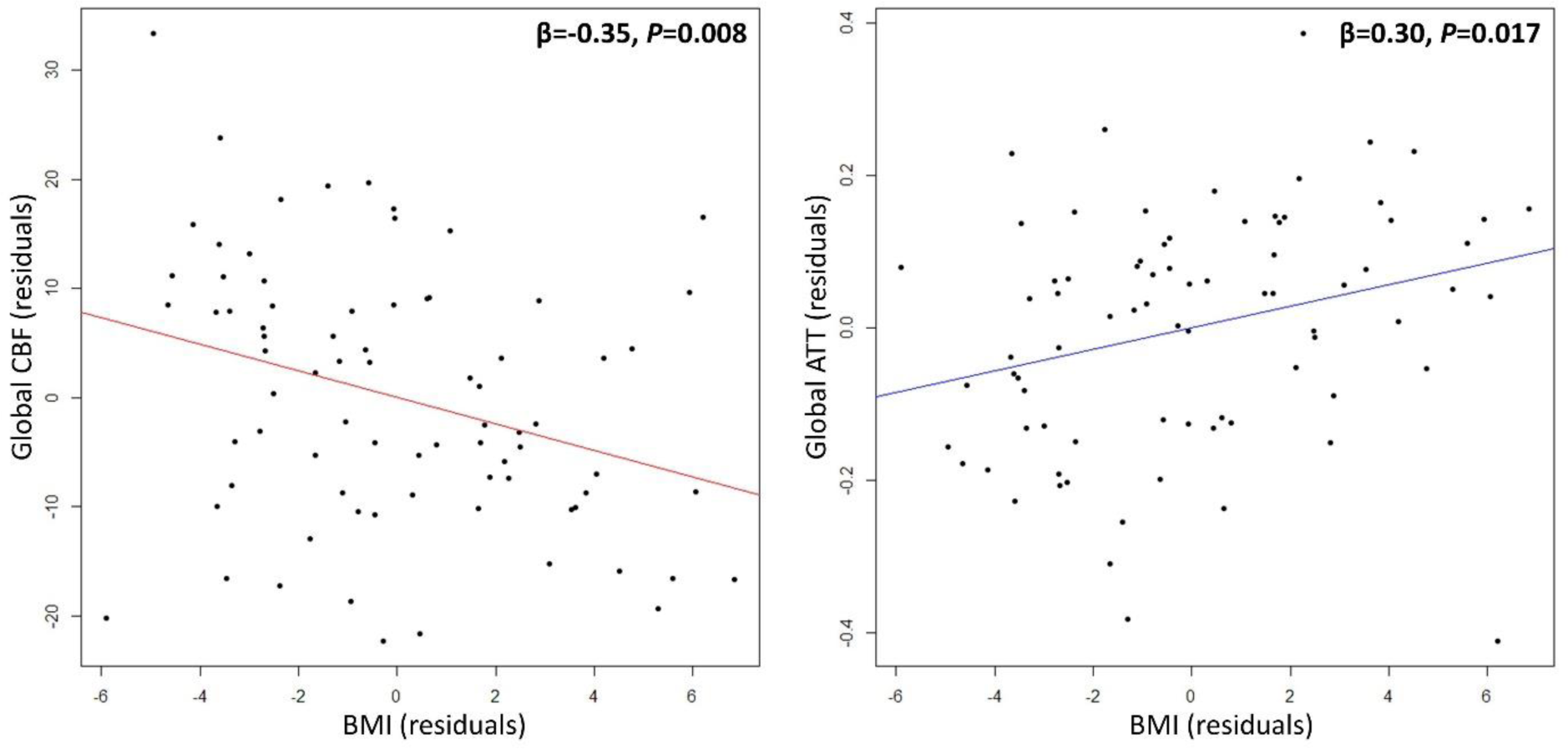
Partial regression association between BMI and global CBF (left) or global ATT (right), adjusted for age, sex, blood pressure, cardiorespiratory fitness, hand grip strength, and grey matter volume. Higher BMI was associated with lower CBF and a longer ATT (n=78). CBF; cerebral blood flow, ATT; arterial transit time, BMI; body mass index.

**Table 2:** Determinants of global resting CBF and ATT in grey matter.

| n=78 | gCBF (mL/100g/min) |  | gATT (s) |  |
| --- | --- | --- | --- | --- |
| | $\beta$ | P | $\beta$ | P |
| Age (years) | -0.16 | 0.298 | 0.43 | **0.004** |
| Sex (1; male, 2; female) | 0.03 | 0.895 | -0.19 | 0.406 |
| BMI (kg/m<sup>2</sup>) | -0.35 | **0.008** | 0.30 | **0.017** |
| SBP (mmHg) | -0.02 | 0.877 | -0.11 | 0.371 |
| DBP (mmHg) | -0.11 | 0.429 | 0.06 | 0.651 |
| $\dot{V}O_{2peak}$ (mL/kg/min) | -0.18 | 0.304 | 0.24 | 0.149 |
| Hand grip strength (kg) | 0.06 | 0.775 | -0.33 | 0.115 |
| Grey matter volume (mm<sup>3</sup>) | 0.06 | 0.687 | -0.01 | 0.943 |

Results from two multiple linear regression analyses. Bold indicates $P < 0.05$ . $\beta$ ; standardised beta coefficient, gCBF; global cerebral blood flow, gATT; global arterial transit time, SBP; systolic blood pressure, DBP; diastolic blood pressure, BMI; body mass index, $\dot{V}O_{2peak}$ ; peak oxygen consumption.

### Determinants of global ATT

The overall global ATT (gATT) model was significant (R^2^adjusted=0.15, F(8,69)=2.66, *P*=0.013). Of the eight independent variables, only age (β=0.43, *P*=0.004; Figure 3) and BMI (β=0.30, *P*=0.017; Figure 2) were significant determinants of gATT, whereby gATT lengthened with both BMI and age. Data shown in Table 2.

**Figure 3:**
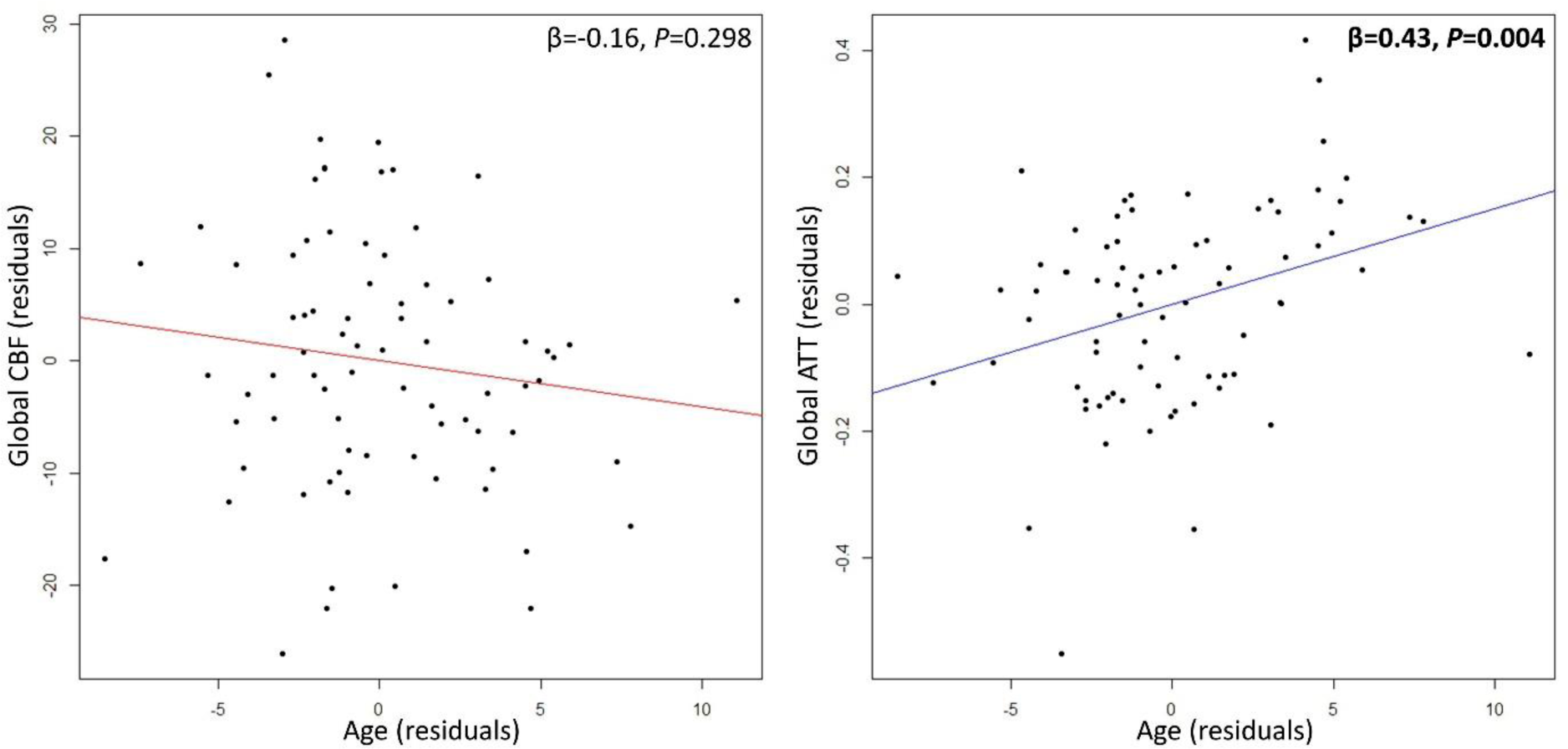
Partial regression association between age and global CBF (left) or global ATT (right), adjusted for sex, BMI, blood pressure, cardiorespiratory fitness, hand grip strength, and grey matter volume. ATT may be more sensitive to age-related changes than CBF (n=78). CBF; cerebral blood flow, ATT; arterial transit time, BMI; body mass index.

### Associations between regional CBF and ATT with age, BMI, and cardiorespiratory fitness

#### Regional CBF

Age and cardiorespiratory fitness were not significantly associated with CBF of any region, before or after adjustment for multiple comparisons. Negative associations between BMI and CBF were present in all regions after adjustment for multiple comparisons, with the largest associations in the temporal (β=-0.44, *P*<0.001), occipital (β=-0.43, *P*<0.001), and parietal (β=-0.41, *P*=0.002) regions.

Data shown in Table S2.

#### Regional ATT

Full regional ATT results are shown in Table 3. Positive associations between age and ATT were present in all regions, only the cingulate region did not survive adjustment for multiple comparisons. The largest association was in the occipital region (β=0.62, *P*<0.001). Significant positive associations between BMI and ATT were present in frontal, parietal, temporal, and motor regions, but these did not survive adjustment for multiple comparisons. There were significant positive associations between cardiorespiratory fitness and ATT in frontal, parietal, occipital and motor regions, associations in parietal (β=0.44, *P*=0.004) and occipital (β=0.45, *P*=0.003) regions survived adjustment for multiple comparisons (Figure 4).

**Figure 4:**
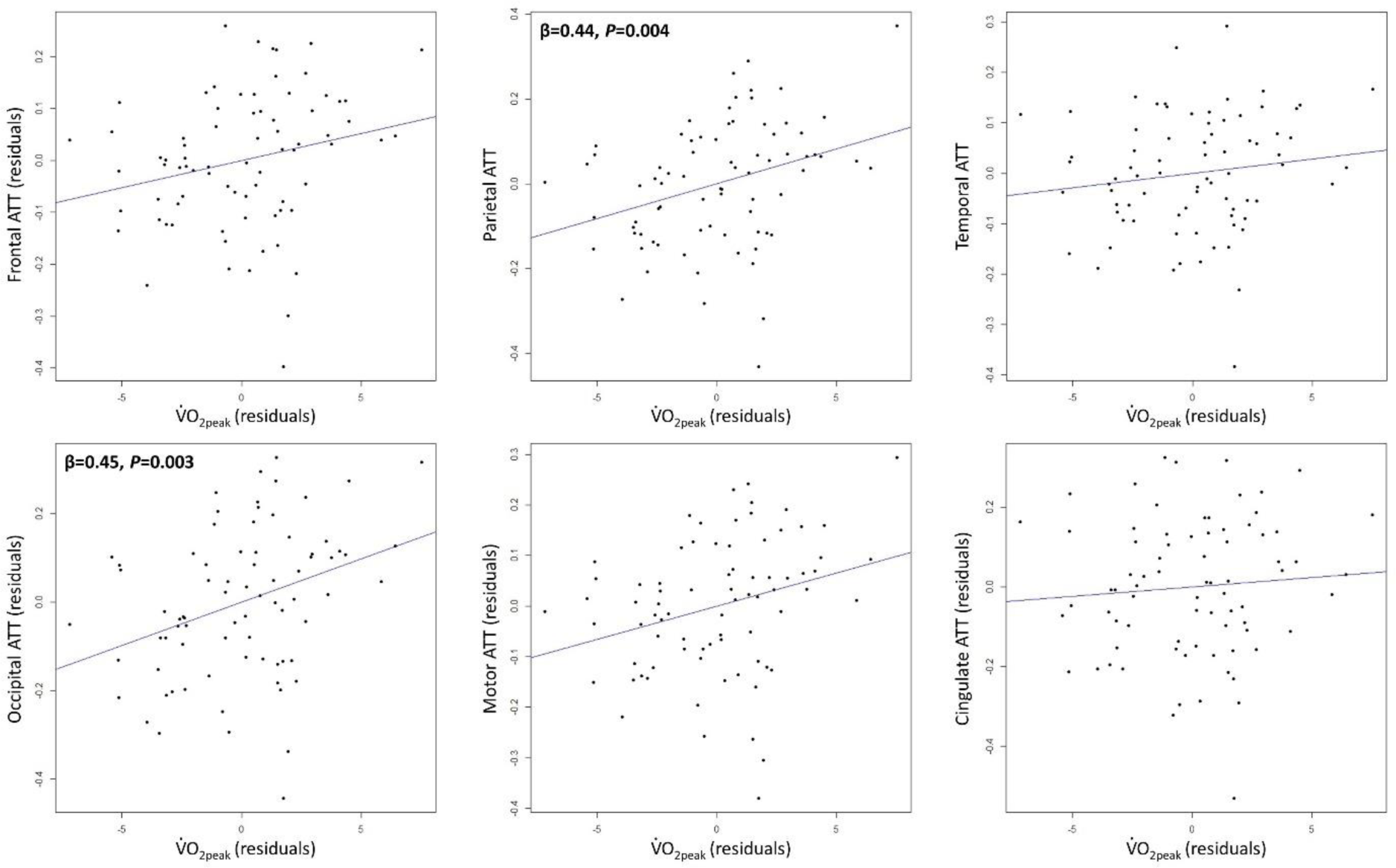
Partial regression associations between regional ATT and cardiorespiratory fitness, adjusted for age, sex, and BMI. Older adults with higher cardiorespiratory fitness experience longer ATT in parietal and temporal regions (n=77). ATT; arterial transit time, BMI; body mass index, *VO*_2*peak*_; peak oxygen consumption.

**Table 3:** Associations between age, BMI, and cardiorespiratory fitness with regional ATT.

| ATT | Age | | BMI | | $\dot{V}O_{2peak}$ | |
| --- | --- | --- | --- | --- | --- | --- |
| | $\beta$ | *P* | $\beta$ | *P* | $\beta$ | *P* |
| Frontal | 0.47 | **<0.001** | 0.24 | 0.046 | 0.33 | 0.042 |
| Parietal | 0.57 | **<0.001** | 0.23 | 0.040 | 0.44 | **0.004** |
| Temporal | 0.46 | **<0.001** | 0.25 | 0.043 | 0.20 | 0.225 |
| Occipital | 0.62 | **<0.001** | 0.19 | 0.083 | 0.45 | **0.003** |
| Motor | 0.49 | **<0.001** | 0.26 | 0.034 | 0.40 | 0.012 |
| Cingulate | 0.32 | 0.017 | 0.24 | 0.063 | 0.12 | 0.495 |

Separate multiple linear regressions were performed for each region ( $n=77$ ), independent variables: age, sex, BMI, and $\dot{V}O_{2peak}$ . Bold indicates significant *P* values survived adjustment for multiple comparisons. $\beta$ ; standardised beta coefficient, ATT; arterial transit time, BMI; body mass index, $\dot{V}O_{2peak}$ ; peak oxygen consumption.

#### Associations between cognitive function and global CBF or ATT

Overall, the models including all independent variables did not significantly explain variance in gCBF (R^2^adjusted=-0.017, F(9,66)=0.86, *P*=0.56) or gATT (R^2^adjusted=0.031, F(9,66)=1.27, *P*=0.27). Measures of processing speed, working memory, or attention were not significantly associated with gCBF or gATT. The only noteworthy result between both models was that processing speed accuracy approached significance for predicting gCBF (β=0.25, *P*=0.053). Global regression data and summarised results from regional analyses can be found in Section 5 of the supplementary material.

## Discussion

The present study aimed to identify modifiable determinants of CBF and ATT in healthy older adults, and assess whether CBF and ATT are associated with cognitive function. CBF calculations were adjusted for differences in ATT, an important, but often ignored, factor to consider, especially in older populations.^60^ The present data show older adults with a higher BMI had lower global CBF and a longer global ATT, and that global ATT lengthened with age. Sex, blood pressure, cardiorespiratory fitness, hand grip strength, or grey matter volume were not significant determinants of global CBF or ATT. Regional analysis confirmed a lack of association between cardiorespiratory fitness and CBF, but indicated that older adults with a higher cardiorespiratory fitness experience longer ATT in parietal and occipital regions. Cognitive function was not associated with CBF or ATT, globally or regionally.

### Higher BMI predicts lower CBF and prolonged ATT in healthy older adults

BMI was the only significant determinant of global CBF, whereby higher BMI was associated with lower CBF (decrease 1.2 mL/100g/min per 1 kg/m^2^ increase). This relationship was robust and was present in all regional analyses. This finding agrees with previous population-based research that report negative associations between global CBF and BMI in older adults.^25,26^ Similarly, compared to healthy older adults, global CBF is 15% lower in those diagnosed with metabolic syndrome.^24^ Furthermore, BMI-associated CBF reductions are greater than that of age, highlighting the importance of managing obesity for the maintenance of brain vascular health in older adults.^25^

A novel finding from the present study was that older adults with a higher BMI have a prolonged global ATT. Although previous research has used ASL to investigate the relationship between CBF and obesity,^24,25^ multiple post-label delays were not used and therefore ATT was not measured. To the best of our knowledge, only one other study has investigated the relationship between BMI and ATT, reporting that a higher BMI was associated with a shorter regional ATT (limited to clusters within the occipital lobe).^65^ Although this contradicts the present data, making a direct comparison is problematic due to differences with the previously studied population (male coronary artery disease patients). Therefore, the present findings reinforce the known damaging effects of excessive weight gain on brain vascular health.

The mechanisms mediating the relationship between BMI and poor cardiovascular health are poorly understood. Nevertheless, the prevalence of metabolic and vascular risk factors in those with a high BMI is common (e.g., hypertension, arterial stiffness, endothelial dysfunction, and hyperlipidaemia).^81,82^ These risk factors are known to worsen cerebrovascular health^83,84^ and are associated with lower CBF.^27,85,86^ Given that the present population was normotensive with only 13 participants (17%) taking lipid-lowering medication, and that blood pressure or grey matter volume were not significant determinants of global CBF or ATT, it is unlikely that the higher blood pressure, hyperlipidaemia, or cerebral atrophy^87^ typically associated with higher BMI explain the observed results. Therefore, adverse structural and functional changes to peripheral and cerebral vessels associated with higher BMI are potentially the mediators of these associations.

Taken together, poor metabolic health and excessive weight gain have deleterious effects on brain vascular health. Fortunately, there are data to suggest that this could be at least partially reversed through modifications to diet and physical activity habits. A one-year weight loss intervention (diet or diet and exercise) in overweight and obese middle-aged adults resulting in ∼10 kg weight loss appears to increase CBF across large regions of the brain.^88^ Evidence also indicates that engagement with high, but not low or moderate, levels of physical activity ameliorates the CBF reductions observed with higher BMI.^25^ The impact of weight loss interventions on ATT warrants further investigation.

### Cardiorespiratory fitness lacks association with CBF, whereas associations with ATT may be region-specific

There are conflicting findings regarding the association between cardiorespiratory fitness and CBF in older adults, and only one previous study using ASL has adjusted CBF for ATT differences.^52^ Fitter older adults have lower blood pressure,^89^ better endothelial function,^90^ more elastic central and cerebral arteries,^91^ and increased grey matter volume.^92^ These factors have been associated with greater CBF.^17,19,27,86^ However, the present data found no association between cardiorespiratory fitness and global or regional CBF. Other cross-sectional ASL studies report no global effect,^33,52,54^ but positive regional associations are generally observed.^33,36–39^ Previously reported regional effects often relate to much smaller regions than those examined in the present study (e.g., occipitoparietal area, posterior cingulate cortex, precuneus, middle frontal gyrus), and CBF was not adjusted for ATT differences in those studies. cardiorespiratory fitness as a large genetic component (∼47%)^32^ and refers primarily to the efficiency of oxygen delivery/utilisation at skeletal muscle, not the brain, potentially explaining the present findings. Accelerometer-measured physical activity could have stronger associations with CBF.^22,23^

Cardiorespiratory fitness was not a significant determinant of global ATT, but higher cardiorespiratory fitness was associated with longer ATT in parietal and occipital regions, with trends also seen in frontal and motor regions. These regional findings oppose the hypothesis that higher cardiorespiratory fitness reduces resistance to blood velocity (e.g., vessel tortuosity, arterial stenosis, collateral recruitment) and thus shortens ATT. The only other study investigating the association between cardiorespiratory fitness and ATT reported no association in older adults (n=14) but regional analysis, using the same ROIs as the present study (except temporal), did show non-significant elevations in ATT of high vs. low fitness older adults.^52^ The present data indicate that, in healthy older adults, cardiorespiratory fitness does not alter the delivery rate of arterial blood to cerebral tissue, but instead lengthens the time taken for blood to arrive at parietal and occipital regions from larger cerebral arteries in the neck.

The cardiorespiratory fitness-related prolongation of regional ATT could be related to the blood velocity, the vascular path length, or both. A faster cerebral artery blood velocity (CBv) is associated with shorter ATT^52^ and is evident with higher cardiorespiratory fitness.^34,40^ However, associations between cardiorespiratory fitness and CBv are now thought to be sex-dependent^56,58^ and a meta-analysis actually reports no significant association (n=886).^93^ This ambiguity suggests changes in large artery blood velocity are not responsible for changes in regional ATT. Lifelong engagement with physical activity markedly attenuates age-related increases in cerebral vessel tortuosity,^94^ suggesting that increased vascular path length is not the cause of prolonged ATT. The same study also demonstrated that physically active older adults have more small cerebral vessels. Given that total vessel cross-sectional area is inversely proportional to blood velocity (assuming constant blood flow), cardiorespiratory fitness-induced small vessel cerebral angiogenesis may be slowing blood velocity and thus prolonging regional ATT. Interestingly, a longer ATT has been associated with greater oxygen extraction fraction on the affected side of stenosis patients,^95^ possibly indicating that the overall slower blood velocities within the cerebral vascular tree also translate into longer capillary transit times, thus improving gas exchange and oxygen extraction fraction. Therefore, older adults with a higher cardiorespiratory fitness may benefit from an improved cerebral oxygen extraction fraction, which is known to be true for skeletal muscle,^96^ and could help explain the preservation of cerebral tissue integrity and cognition that is associated with regular exercise training.

### Age was not a significant determinant of CBF, but was associated with prolonged ATT

Age-related CBF decline is well documented^4,18,19^ but was not replicated in the present data (although a trend was observed). This is likely due to the limited age range (60–81 yrs), with 56% of participants aged ≤65 and only 17% aged ≥70. Age-related cerebral atrophy and/or reductions in cerebral metabolic rate are hypothesised to explain CBF declines.^97^ However, regional CBF reductions can be independent of atrophy^98^ and a bidirectional relationship exists between CBF and brain volume.^99^ Furthermore, although overall cerebral metabolic rate declines with age, this is due to atrophy, and the metabolic rate of remaining tissue actually increases with age.^100^ Systemic age-related cardiovascular and cerebrovascular deterioration likely contribute to CBF reductions. For example, hypertension, arterial stiffness, and endothelial dysfunction typically increase with age,^101,102^ and have all been associated with lower resting CBF.^19,27,86,100^

Age was however a significant determinant of global and regional ATT in the present study, conforming with pervious findings.^4,62^ The present data indicates global ATT slows by 150 ms per decade in healthy older adults, with occipital and parietal regions experiencing the greatest effects. The fact that both age and cardiorespiratory fitness are associated with a longer ATT in older adults is somewhat contradictive given that this is generally considered a marker of worsening cerebrovascular health. However, the cause of prolonged ATT likely differs between age and cardiorespiratory fitness. As discussed, cardiorespiratory fitness-induced ATT prolongation may be due to cerebral angiogenesis increasing the number of small cerebral vessels and thus reducing blood velocity within this region of the cerebral vascular tree. Conversely, adverse structural changes to the cerebrovasculature, such as increases in cerebral vessel tortuosity^103,104^ and the prevalence of cerebral stenosis,^105^ may be the primary drivers of age-related increases in ATT. Furthermore, blood velocity within larger cerebral vessels is well documented to decline with age.^34,56^ Although age-related ATT prolongation occurs due to cerebrovascular deterioration, it may consequently serve to improve oxygen extraction fraction (similarly to cardiorespiratory fitness) and help explain why this and the cerebral metabolic rate of remaining cerebral tissue increases with age.^100^ However, cardiorespiratory fitness-related ATT prolongation should be considered a marker of beneficial cerebrovascular adaptation.

Given that age was a significant determinant of global and regional ATT but not CBF (despite being well reported in the literature), ATT may be more sensitive to age-related decline (Figure 3).

Therefore, changes in ATT could have some prognostic value identifying the onset of cerebrovascular impairment in healthy older adults with low cardiorespiratory fitness.

### Blood pressure was not a determinant of CBF or ATT

The present study found no association between global CBF or ATT and blood pressure in healthy older adults (without hypertension). Previous population-based research found higher blood pressure was associated with lower global CBF in older adults (n=468, 34% hypertensive).^19^ Differences in the hypertensive status of the sample or the ASL protocol used may explain discrepancies with the present data. However, other research reports no association between blood pressure and global CBF either cross-sectionally or after ∼3 yr follow-up in hypertensive older adults, but those using anti-hypertensive medication did have lower CBF.^106^ The relationship between blood pressure and CBF is clearly complex. For example, hypertension is actually suggested to be a protective response to cerebral hypoperfusion in attempt to maintain CBF.^107^ Therefore, between-study differences may be explained by variance in the severity or duration of blood pressure changes experienced.

The present study is believed to be the first investigating associations between ATT and blood pressure in older adults. Higher blood pressure was hypothesised to be associated with longer ATT as this can induce damage to the cerebral tissue and vasculature.^108–110^ However, theoretically, the inverse could also be true. Higher blood pressures increase cerebral perfusion pressure, potentially accelerating blood velocity and thus shortening ATT. Likewise to CBF, the duration of exposure to high blood pressures is likely an important factor when considering its impact on ATT (i.e., ATT may shorten in the short-term but lengthen in the long-term).

Alternative multiple linear regressions were performed with either mean arterial pressure or pulse pressure but this had no effect on results (data not shown). Therefore, the present study indicates that blood pressure is not a significant determinant of global CBF or ATT in healthy older adults without hypertension. Conclusions may differ in hypertensive populations or longitudinal investigations. Given that higher blood pressure increases the risk of cerebrovascular dysfunction,^111^ cardiovascular diseases,^112^ and dementia,^113^ it can be assumed that maintaining blood pressure within the normal healthy range is beneficial for brain health in older adults.

### No association between CBF or ATT and cognitive function

Chronic cerebral hypoperfusion, resulting in limited energy availability of cerebral tissue, is emerging as a potential key factor of cognitive decline in older adults.^114^ However, the present data found no association between global or regional CBF with processing speed, working memory, or attention in healthy older adults. This agrees with previous findings that global and regional CBF are not associated with global cognition, memory, attention, or executive functioning in cognitively normal older adults (n=113).^16^ Interestingly, however, a follow-on study to this aforementioned study did find longitudinal associations between CBF and cognitive function in a subset of these participants (n=89) (2 yr follow-up).^14^ Specifically, a lower baseline CBF (global, frontal, parietal, temporal, and occipital regions) predicted a greater decline in global cognition and attention/psychomotor speed whereas only frontal and temporal baseline CBF predicted decline in memory, and that baseline global or regional CBF were not predictive of executive or language functions. Collectively, these data do suggest an importance of CBF for cognitive function, but this is both domain- and region-specific, and only apparent when analysed over time. This temporal association between CBF and cognitive function has been reported elsewhere.^10^

The present study found no associations between global or regional ATT and cognitive function in healthy older adults. To the best of our knowledge, this is the first study to examine these relationships. Previous research have only investigated relationships between ATT and clinical cognitive impairment, reporting that ATT is prolonged in Alzheimer’s disease and this was associated with lower mini–mental state examination (MMSE) scores in two out of ten regions studied.^3^

It appears that global or regional CBF or ATT are not associated with contemporaneous cognitive function in healthy older adults, but they may still help explain or predict changes in cognitive function over time. The predictive capacity of CBF has been shown previously,^14^ although understanding of which regions and cognitive domains are most affected by could be improved. Given that the present data indicates that ATT may be more sensitive to age-related decline than CBF, the predictive capacity of ATT should be investigated as it could be an early biomarker of cerebrovascular-related cognitive decline in healthy populations. Alternatively, resting state cerebrovascular haemodynamics may not be key determinants of cognitive function. Instead of oxygen delivery, oxygen utilisation of the cerebral tissue and thus neuronal function may have a larger impact.

## Future Directions

This cross-sectional analysis does not account for variations in genetics or pre-existing cardiovascular, metabolic, and cerebrovascular health. Longitudinal research is needed to make robust conclusions. There are likely other more influential determinants of CBF not investigated in the present study, such as physical activity, arterial stiffness, endothelial function, cerebrovascular reactivity, cerebral metabolic rate, blood lipids, and social activities that deserve future investigation. Regarding cognition, future research should not only use contemporaneous measurements, but investigate whether CBF and ATT are reliable predictors of cognitive decline over time. CBF and ATT are markers of cerebrovascular health, but more general brain health may be more strongly dictated by the ability of the cerebral tissue to extract and use essential nutrients delivered in the blood (i.e., oxygen extraction fraction and cerebral metabolic rate). Future research should investigate associations between these variables with cardiorespiratory fitness, CBF, ATT, and cognitive function. The present study lacked dietary controls (e.g., no restriction on caffeine or polyphenols) prior to MRI acquisition, which could have impacted results.^115–117^ Similarly, participants’ COVID-19 infection history was not known, thought to reduce CBF,^118^ but all were vaccinated and free from symptoms during testing. The most powerful regulator of CBF is arterial partial pressure of CO2 (*P*aCO2).^119^ *P*aCO2 or its proxy, partial pressure of end-tidal CO2 (*P*etCO2),^120^ was not measured during CBF/ATT measurements, but can be easily manipulated by ventilation changes that could occur during an MRI scan. Future studies should measure and correct for *P*etCO2.

## Conclusion

This study aimed to identify modifiable determinants of CBF and ATT in healthy older adults and assess whether these markers of brain vascular health were associated with cognitive function. Results show that older adults with a higher BMI have lower global CBF and a longer global ATT. Global ATT lengthened with age, whereas no association was found with CBF, possibly indicating a greater sensitivity of ATT to age-related decline. cardiorespiratory fitness was not a determinant of global CBF or ATT, but regional analysis unexpectedly found that cardiorespiratory fitness was associated with a longer ATT in parietal and occipital regions. The responses of CBF and ATT to exercise training warrant further investigation as longitudinal research reduces the obscuring effects of pre-existing health status and lifestyle habits on results. Regarding cognitive function, global or regional CBF or ATT were not associated with processing speed, working memory, or attention; however, future research should instead longitudinally investigate the predictive capacity of these brain vascular health markers.

## Acknowledgements

This work was funded by the Norwegian Research Council (FRIPRO 300030). We thank Bethany Skinner, Consuelo Vidal Gran, Nicolas Hayston, Rupali Limachya, Amelie Grandjean, Aoife Marley, Shi Miao, and Samuel Thomas for data collection support, and Roksana Markiewicz for cognitive data analysis. We thank Danny Wang and the University of Southern California’s Steven Neuroimaging and Informatics Institute for the provision of the pCASL sequence used in this work, which was provided through a C2P agreement with The Regents of the University of California.

## Author contributions

- Conceptualisation: JF, KS, SJEL, KJM.
- Implementation: JF, KS, SJEL, KJM, SHF, FR, KEJ, AG, SB, HL.
- Data collection: JF, FR, KEJ.
- Data analysis: JF, KJM.
- Data visualisation: JF.
- Writing (original draft): JF, KS, SJEL.
- Writing (reviewing and editing): KF, KS, SJEL, KJM, FR, KEJ, SHF, SB, HL.

## Authors

Jack Feron, Katrien Segaert, Foyzul Rahman, Sindre H Fosstveit, Kelsey E Joyce, Ahmed Gilani, Hilde Lohne-Seiler, Sveinung Berntsen, Karen J Mullinger, Samuel J E Lucas

## Supplementary material

Supplementary material contains additional information regarding Materials and Methods and Results sections.

## References

1. United Nations. World Population Prospects 2022: Summary of Results. *Department of Economic and Social Affairs* 2022; UN DESA/POP/2021/TR/NO. 3.

2. Wolters, F. J. et al. Cerebral Perfusion and the Risk of Dementia: A Population-Based Study. Circulation 2017; 136: 719–728.

3. Sun, M. et al. Potential Diagnostic Applications of Multi-Delay Arterial Spin Labeling in Early Alzheimer’s Disease: The Chinese Imaging, Biomarkers, and Lifestyle Study. Front. Neurosci 2022; 16: 934471.

4. Damestani, N. L. et al. Associations between age, sex, APOE genotype, and regional vascular physiology in typically aging adults. NeuroImage 2023; 275: 120167.

5. Salthouse, T. Consequences of Age-Related Cognitive Declines. Annu. Rev. Psychol 2012; 63: 201–226.

6. Stites, S. D., Harkins, K., Rubright, J. D. & Karlawish, J. Relationships Between Cognitive Complaints and Quality of Life in Older Adults With Mild Cognitive Impairment, Mild Alzheimer Disease Dementia, and Normal Cognition. Alzheimer Disease & Associated Disorders 2018; 32: 276–283.

7. Salthouse, T. Selective review of cognitive aging. J Int Neuropsychol Soc 2010; 16: 754–760.

8. Lee, R. H. C. et al. Cerebral ischemia and neuroregeneration. Neural Regen Res 2018; 13: 373.

9. Love, S. & Miners, J. S. Cerebrovascular disease in ageing and Alzheimer’s disease. Acta Neuropathol 2016; 131: 645–658.

10. De Vis, J. B. et al. Arterial-spin-labeling (ASL) perfusion MRI predicts cognitive function in elderly individuals: A 4-year longitudinal study. Magnetic Resonance Imaging 2018; 48: 449– 458.

11. Ebenau, J. L. et al. Cerebral blood flow, amyloid burden, and cognition in cognitively normal individuals. Eur J Nucl Med Mol Imaging 2023; 50: 410–422.

12. Leeuwis, A. E. et al. Cerebral Blood Flow and Cognitive Functioning in a Community-Based, Multi-Ethnic Cohort: The SABRE Study. Front. Aging Neurosci 2018; 10: 279.

13. Moonen, J. E. et al. Contributions of Cerebral Blood Flow to Associations Between Blood Pressure Levels and Cognition: The Age, Gene/Environment Susceptibility-Reykjavik Study. Hypertension 2021; 77: 2075–2083.

14. van Dinther, M. et al. Lower cerebral blood flow predicts cognitive decline in patients with vascular cognitive impairment. Alzheimer’s & Dementia 2023. doi:10.1002/alz.13408.

15. Hshieh, T. T. et al. Cerebral blood flow MRI in the nondemented elderly is not predictive of post-operative delirium but is correlated with cognitive performance. J Cereb Blood Flow Metab 2017; 37: 1386–1397.

16. Leeuwis, A. E., et al. Cerebral blood flow and cognitive functioning in patients with disorders along the heart–brain axis: Cerebral blood flow and the heart–brain axis. A&D Transl Res & Clin Interv 2020; 6: e12034.

17. Poels, M. M. et al. Total Cerebral Blood Flow in Relation to Cognitive Function: The Rotterdam Scan Study. J Cereb Blood Flow Metab 2008: 28; 1652–1655.

18. Dijsselhof, M. B. J. et al. The value of arterial spin labelling perfusion MRI in brain age prediction. Human Brain Mapping 2023; 44; 2754–2766.

19. Leidhin, C. N. et al. Age-related normative changes in cerebral perfusion: Data from The Irish Longitudinal Study on Ageing (TILDA). NeuroImage 2021; 229; 117741.

20. Bangen, K. J. et al. Greater accelerometer-measured physical activity is associated with better cognition and cerebrovascular health in older adults. J Int Neuropsychol Soc 2023; 29: 859–869.

21. Boraxbekk, C.-J., Salami, A., Wåhlin, A. & Nyberg, L. Physical activity over a decade modifies age-related decline in perfusion, gray matter volume, and functional connectivity of the posterior default-mode network—A multimodal approach. NeuroImage 2016; 131: 133–141.

22. Zlatar, Z. Z. et al. Dose-dependent association of accelerometer-measured physical activity and sedentary time with brain perfusion in aging. Experimental Gerontology 2019; 125: 110679.

23. Sanders, A. et al. Associations between everyday activities and arterial spin labeling-derived cerebral blood flow: A longitudinal study in community-dwelling elderly volunteers. Human Brain Mapping 2023; 44; 3377–3393.

24. Birdsill, A. C. et al. Low cerebral blood flow is associated with lower memory function in metabolic syndrome. Obesity 2013; 21: 1313–1320.

25. Knight, S. P. et al. Obesity is associated with reduced cerebral blood flow – modified by physical activity. Neurobiology of Aging 2021; 105; 35–47.

26. Vernooij, M. W. et al. Total Cerebral Blood Flow and Total Brain Perfusion in the General Population: The Rotterdam Scan Study. J Cereb Blood Flow Metab 2008; 28: 412–419.

27. Jefferson, A. L. et al. Higher Aortic Stiffness Is Related to Lower Cerebral Blood Flow and Preserved Cerebrovascular Reactivity in Older Adults. Circulation 2018; 138: 1951–1962.

28. Tomoto, T. et al. One-year aerobic exercise increases cerebral blood flow in cognitively normal older adults. J Cereb Blood Flow Metab 2023; 43: 404–418.

29. Colcombe, S. & Kramer, A. F. Fitness Effects on the Cognitive Function of Older Adults: A Meta-Analytic Study. Psychol Sci 2003; 14: 125–130.

30. Ludyga, S., Gerber, M., Pühse, U., Looser, V. N. & Kamijo, K. Systematic review and meta-analysis investigating moderators of long-term effects of exercise on cognition in healthy individuals. Nat Hum Behav 202; 4: 603–612.

31. Northey, J. M., Cherbuin, N., Pumpa, K. L., Smee, D. J. & Rattray, B. Exercise interventions for cognitive function in adults older than 50: a systematic review with meta-analysis. Br J Sports Med 2018; 52: 154–160.

32. Bouchard, C. et al. Familial aggregation of VO_2_ max response to exercise training: results from the HERITAGE Family Study. Journal of Applied Physiology 1999; 87: 1003–1008.

33. Thomas, B. P. et al. Life-long aerobic exercise preserved baseline cerebral blood flow but reduced vascular reactivity to CO 2. Magnetic Resonance Imaging 2013; 38: 1177–1183.

34. Ainslie, P. N. et al. Elevation in cerebral blood flow velocity with aerobic fitness throughout healthy human ageing. The Journal of Physiology 2008; 586: 4005–4010.

35. Willie, C. K. et al. Utility of transcranial Doppler ultrasound for the integrative assessment of cerebrovascular function. Journal of Neuroscience Methods 2011; 196: 221–237.

36. Dougherty, R. J. et al. Fitness, independent of physical activity is associated with cerebral blood flow in adults at risk for Alzheimer’s disease. Brain Imaging and Behavior 2023; 14: 1154–1163.

37. Johnson, N. F. et al. Cardiorespiratory fitness modifies the relationship between myocardial function and cerebral blood flow in older adults. NeuroImage 2016; 131: 126–132.

38. Tarumi, T. et al. Central artery stiffness, neuropsychological function, and cerebral perfusion in sedentary and endurance-trained middle-aged adults. Journal of Hypertension 2013; 31: 2400– 2409.

39. Zimmerman, B. et al. Cardiorespiratory fitness mediates the effects of aging on cerebral blood flow. Front. Aging Neurosci. 2014; 6.

40. Bailey, D. M. et al. Elevated Aerobic Fitness Sustained Throughout the Adult Lifespan Is Associated With Improved Cerebral Hemodynamics. Stroke 2013; 44: 3235–3238.

41. Alfini, A. J. et al. Hippocampal and Cerebral Blood Flow after Exercise Cessation in Master Athletes. Front. Aging Neurosci 2016; 8.

42. Alfini, A. J., Weiss, L. R., Nielson, K. A., Verber, M. D. & Smith, J. C. Resting Cerebral Blood Flow After Exercise Training in Mild Cognitive Impairment. JAD 2019; 6:, 671–684.

43. Chapman, S. B. et al. Shorter term aerobic exercise improves brain, cognition, and cardiovascular fitness in aging. Front. Aging Neurosci. 2013; 5.

44. Kleinloog, J. P. D. et al. Aerobic Exercise Training Improves Cerebral Blood Flow and Executive Function: A Randomized, Controlled Cross-Over Trial in Sedentary Older Men. Front. Aging Neurosci. 2019; 11: 333.

45. Moore, S. A. et al. Effects of Community Exercise Therapy on Metabolic, Brain, Physical, and Cognitive Function Following Stroke: A Randomized Controlled Pilot Trial. Neurorehabil Neural Repair 2015; 29: 623–635.

46. Akazawa, N., et al. Aerobic exercise training increases cerebral blood flow in postmenopausal women. ARTRES 2012: 6; 124.

47. Bailey, T. G. et al. Exercise training reduces the frequency of menopausal hot flushes by improving thermoregulatory control. Menopause 2016; 23: 708–718.

48. Guadagni, V. et al. Aerobic exercise improves cognition and cerebrovascular regulation in older adults. Neurology 2023; 94: e2245–e2257.

49. Murrell, C. J. et al. Cerebral blood flow and cerebrovascular reactivity at rest and during sub- maximal exercise: Effect of age and 12-week exercise training. AGE 2013; 35: 905–920.

50. Intzandt, B. et al. Higher cardiovascular fitness level is associated with lower cerebrovascular reactivity and perfusion in healthy older adults. J Cereb Blood Flow Metab 2020; 40: 1468– 1481.

51. Olivo, G. et al. Higher VO2max is associated with thicker cortex and lower grey matter blood flow in older adults. Sci Rep 2021; 11: 16724.

52. Burley, C. V., Francis, S. T., Whittaker, A. C., Mullinger, K. J. & Lucas, S. J. E. Measuring resting cerebral haemodynamics using MRI arterial spin labelling and transcranial Doppler ultrasound: Comparison in younger and older adults. Brain and Behavior 2021; 11: e02126.

53. Flodin, P., Jonasson, L. S., Riklund, K., Nyberg, L. & Boraxbekk, C. J. Does Aerobic Exercise Influence Intrinsic Brain Activity? An Aerobic Exercise Intervention among Healthy Old Adults. Front. Aging Neurosci 2017; 9: 267.

54. Krishnamurthy, V. et al. The Relationship Between Resting Cerebral Blood Flow, Neurometabolites, Cardio-Respiratory Fitness and Aging-Related Cognitive Decline. Front. Psychiatry 2023; 13: 923076.

55. Flück, D. et al. Age, aerobic fitness, and cerebral perfusion during exercise: role of carbon dioxide. American Journal of Physiology-Heart and Circulatory Physiology 2014; 307: H515– H523.

56. Lefferts, W. K. et al. Influence of sex and presence of cardiovascular risk factors on relations between cardiorespiratory fitness and cerebrovascular hemodynamics. Journal of Applied Physiology 2022; 133: 1019–1030.

57. Vicente-Campos, D. et al. Impact of a physical activity program on cerebral vasoreactivity in sedentary elderly people. J Sports Med Phys Fitness 2012; 52: 537–544.

58. Zeller, N. P. et al. Sex-specific effects of cardiorespiratory fitness on age-related differences in cerebral hemodynamics. Journal of Applied Physiology 2022; 132: 1310–1317.

59. Zhu, Y.-S. et al. Cerebral Vasomotor Reactivity during Hypo- and Hypercapnia in Sedentary Elderly and Masters Athletes. J Cereb Blood Flow Metab 2013; 33: 1190–1196.

60. Dai, W. et al. Effects of arterial transit delay on cerebral blood flow quantification using arterial spin labeling in an elderly cohort: Arterial Transit Delay Effect on Elderly CBF. J. Magn. Reson. Imaging 2017; 45: 472–481.

61. Liu, Y. et al. Arterial spin labeling MRI study of age and gender effects on brain perfusion hemodynamics. Magnetic Resonance in Med 2012; 68: 912–922.

62. Mutsaerts, H. J. M. M. et al. Cerebral Perfusion Measurements in Elderly with Hypertension Using Arterial Spin Labeling. PLoS ONE 2015; 10: e0133717.

63. Yu, H. et al. Enhanced Arterial Spin Labeling Magnetic Resonance Imaging of Cerebral Blood Flow of the Anterior and Posterior Circulations in Patients With Intracranial Atherosclerotic Stenosis. Front. Neurosci. 2022; 15: 823876.

64. Mak, H. K. F. et al. Quantitative Assessment of Cerebral Hemodynamic Parameters by QUASAR Arterial Spin Labeling in Alzheimer’s Disease and Cognitively Normal Elderly Adults at 3-Tesla. JAD 2012; 31: 33–44.

65. MacIntosh, B. J. et al. Regional Cerebral Arterial Transit Time Hemodynamics Correlate with Vascular Risk Factors and Cognitive Function in Men with Coronary Artery Disease. AJNR Am J Neuroradiol 2015; 36: 295–301.

66. Bull, F. C. et al. World Health Organization 2020 guidelines on physical activity and sedentary behaviour. Br J Sports Med 2020; 54: 1451–1462.

67. Nasreddine, Z. S. et al. The Montreal Cognitive Assessment, MoCA: A Brief Screening Tool For Mild Cognitive Impairment. J American Geriatrics Society 2005; 53: 695–699.

68. Borg, G. A. Psychophysical bases of perceived exertion. Med Sci Sports Exerc 1982; 14: 377– 381.

69. Mamen, A., Laparidist, C. & van den Tillaar, R. Precision in Estimating Maximal Lactate Steady State Performance in Running Using a Fixed Blood Lactate Concentration or a Delta Value from an Incremental Lactate Profile Test. nternational Journal of Applied Sports Sciences 2011; 23: 212–224.

70. Rahman, F. et al. Lifestyle and brain health determinants of word-finding failures in healthy ageing. 2023. Preprint at 10.1101/2023.12.08.570799.

71. Kilroy, E. et al. Reliability of two-dimensional and three-dimensional pseudo-continuous arterial spin labeling perfusion MRI in elderly populations: Comparison with 15o-water positron emission tomography. Magnetic Resonance Imaging 2014; 39: 931–939.

72. Wang, D. J. J. et al. Multi-delay multi-parametric arterial spin-labeled perfusion MRI in acute ischemic stroke — Comparison with dynamic susceptibility contrast enhanced perfusion imaging. NeuroImage: Clinical 2013; 3: 1–7.

73. Fan, A. P., Jahanian, H., Holdsworth, S. J. & Zaharchuk, G. Comparison of cerebral blood flow measurement with [^15^ O]-water positron emission tomography and arterial spin labeling magnetic resonance imaging: A systematic review. J Cereb Blood Flow Metab 2016; 36: 842– 861.

74. Woods, J. G., Chappell, M. A. & Okell, T. W. A general framework for optimizing arterial spin labeling MRI experiments. Magnetic Resonance in Med 2019; 81: 2474–2488.

75. Chappell, M. A., Groves, A. R., Whitcher, B. & Woolrich, M. W. Variational Bayesian Inference for a Nonlinear Forward Model. IEEE Trans. Signal Process. 2009; 57: 223–236.

76. Groves, A. R., Chappell, M. A. & Woolrich, M. W. Combined spatial and non-spatial prior for inference on MRI time-series. NeuroImage 2009; 45: 795–809.

77. Woolrich, M. W., Chiarelli, P., Gallichan, D., Perthen, J. & Liu, T. T. Bayesian inference of hemodynamic changes in functional arterial spin labeling data. Magnetic Resonance in Med 2006; 56: 891–906.

78. Chappell, M. A. et al. Partial volume correction of multiple inversion time arterial spin labeling MRI data. Magnetic Resonance in Med 2011; 65: 1173–1183.

79. Smith, S. M. Fast robust automated brain extraction. Human Brain Mapping 2002; 17: 143– 155.

80. Zhang, Y., Brady, M. & Smith, S. Segmentation of brain MR images through a hidden Markov random field model and the expectation-maximization algorithm. IEEE Trans. Med. Imaging 2002; 20: 45–57.

81. Li, P., Wang, L. & Liu, C. Overweightness, obesity and arterial stiffness in healthy subjects: a systematic review and meta-analysis of literature studies. Postgraduate Medicine 2017; 129: 224–230.

82. Tang, N. et al. The effects of the interaction between BMI and dyslipidemia on hypertension in adults. Sci Rep 2022; 12: 927.

83. Pires, P. W., Dams Ramos, C. M., Matin, N. & Dorrance, A. M. The effects of hypertension on the cerebral circulation. American Journal of Physiology-Heart and Circulatory Physiology 2013; 304: H1598–H1614.

84. Suri, M. F. K. et al. Prevalence of Intracranial Atherosclerotic Stenosis Using High-Resolution Magnetic Resonance Angiography in the General Population: The Atherosclerosis Risk in Communities Study. Stroke 2016; 47: 1187–1193.

85. Jennings, J. R. et al. Use of Total Cerebral Blood Flow as an Imaging Biomarker of Known Cardiovascular Risks. Stroke 2013; 44: 2480–2485.

86. Sabayan, B. et al. Markers of endothelial dysfunction and cerebral blood flow in older adults. Neurobiology of Aging 2014; 35: 373–377.

87. Hamer, M. & Batty, G. D. Association of body mass index and waist-to-hip ratio with brain structure: UK Biobank study. Neurology 2019; 92: e594–e600.

88. Stillman, C. M. et al. Changes in cerebral perfusion following a 12-month exercise and diet intervention. Psychophysiology 2021; 58: e13589.

89. Cornelissen, V. A. & Smart, N. A. Exercise Training for Blood Pressure: A Systematic Review and Meta-analysis. J Am Heart Assoc 2013; 2: e004473.

90. Early, K. S. et al. The Effects of Exercise Training on Brachial Artery Flow-Mediated Dilation: A Meta-analysis. Journal of Cardiopulmonary Rehabilitation and Prevention 2017; 37: 77.

91. Shibata, S. et al. The effect of lifelong exercise frequency on arterial stiffness. The Journal of Physiology 2018; 596: 2783–2795.

92. Intzandt, B. et al. Comparing the effect of cognitive vs. exercise training on brain MRI outcomes in healthy older adults: A systematic review. Neuroscience & Biobehavioral Reviews 2021; 128: 511–533.

93. Smith, E. C. et al. Effects of cardiorespiratory fitness and exercise training on cerebrovascular blood flow and reactivity: a systematic review with meta-analyses. American Journal of Physiology-Heart and Circulatory Physiology 2021; 321: H59–H76.

94. Bullitt, E. et al. The Effect of Exercise on the Cerebral Vasculature of Healthy Aged Subjects as Visualized by MR Angiography. AJNR Am J Neuroradiol 2009; 30: 1857–1863.

95. Takeuchi, K. et al. The Utility of Arterial Transit Time Measurement for Evaluating the Hemodynamic Perfusion State of Patients with Chronic Cerebrovascular Stenosis or Occlusive Disease: Correlative Study between MR Imaging and 15O-labeled H2O Positron Emission Tomography. Magn Reson Med Sci 2022; 22: 289–300.

96. Kalliokoski, K. K. et al. Enhanced oxygen extraction and reduced flow heterogeneity in exercising muscle in endurance-trained men. Am J Physiol Endocrinol Metab 2001; 280: E1015–1021.

97. Mokhber, N. et al. Cerebral blood flow changes during aging process and in cognitive disorders: A review. Neuroradiol J 2021; 34: 300–307.

98. Chen, J. J., Rosas, H. D. & Salat, D. H. Age-associated reductions in cerebral blood flow are independent from regional atrophy. NeuroImage 2011; 55: 468–478.

99. Zonneveld, H. I. et al. The bidirectional association between reduced cerebral blood flow and brain atrophy in the general population. J Cereb Blood Flow Metab 2015; 35: 1882–1887.

100. Lu, H. et al. Alterations in Cerebral Metabolic Rate and Blood Supply across the Adult Lifespan. Cerebral Cortex 2011; 21: 1426–1434.

101. Thijssen, D. H. J., Carter, S. E. & Green, D. J. Arterial structure and function in vascular ageing: are you as old as your arteries? The Journal of Physiology 2016; 594: 2275–2284.

102. Toth, P., Tarantini, S., Csiszar, A. & Ungvari, Z. Functional vascular contributions to cognitive impairment and dementia: mechanisms and consequences of cerebral autoregulatory dysfunction, endothelial impairment, and neurovascular uncoupling in aging. Am J Physiol Heart Circ Physiol 2017; 312: H1–H20.

103. Bullitt, E. et al. The effects of healthy aging on intracerebral blood vessels visualized by magnetic resonance angiography. Neurobiology of Aging 2010; 31: 290–300.

104. Chen, L. et al. Quantitative Assessment of the Intracranial Vasculature in an Older Adult Population using iCafe (intraCranial Artery Feature Extraction). Neurobiol Aging 2019; 79: 59– 65.

105. de Weerd, M. et al. Prevalence of Asymptomatic Carotid Artery Stenosis in the General Population. Stroke 2010; 41: 1294–1297.

106. van Dalen, J. W. et al. Longitudinal relation between blood pressure, antihypertensive use and cerebral blood flow, using arterial spin labelling MRI. J Cereb Blood Flow Metab 2021; 41: 1756–1766.

107. Warnert, E. A. H. et al. Is High Blood Pressure Self-Protection for the Brain? Circulation Research 2016; 119: e140–e151.

108. de Roos, A., van der Grond, J., Mitchell, G. & Westenberg, J. Magnetic Resonance Imaging of Cardiovascular Function and the Brain—Is dementia a cardiovascular-driven disease? Circulation 2017; 135: 2178–2195.

109. Jochemsen, H. M. et al. Blood Pressure and Progression of Brain Atrophy: The SMART-MR Study. JAMA Neurology 2013; 70: 1046–1053.

110. Willeit, J. & Kiechl, S. Prevalence and risk factors of asymptomatic extracranial carotid artery atherosclerosis. A population-based study. Arteriosclerosis and Thrombosis: A Journal of Vascular Biology 1993: 13; 661–668.

111. Santisteban, M. M., Iadecola, C. & Carnevale, D. Hypertension, Neurovascular Dysfunction, and Cognitive Impairment. Hypertension 2023; 80: 22–34.

112. Fuchs, F. D. & Whelton, P. K. High Blood Pressure and Cardiovascular Disease. Hypertension 2020; 75: 285–292.

113. Walker, K. A. et al. Association of Midlife to Late-Life Blood Pressure Patterns With Incident Dementia. JAMA 2019; 322: 535–545.

114. Rajeev, V. et al. Chronic cerebral hypoperfusion: a critical feature in unravelling the etiology of vascular cognitive impairment. Acta Neuropathologica Communications 2023; 11: 93.

115. Francis, S. T., Head, K., Morris, P. G. & Macdonald, I. A. The Effect of Flavanol-rich Cocoa on the fMRI Response to a Cognitive Task in Healthy Young People. Journal of Cardiovascular Pharmacology 2006; 47: S215.

116. Lamport, D. J. et al. The effect of flavanol-rich cocoa on cerebral perfusion in healthy older adults during conscious resting state: a placebo controlled, crossover, acute trial. Psychopharmacology (Berl) 2015; 232: 3227–3234.

117. Vidyasagar, R., Greyling, A., Draijer, R., Corfield, D. R. & Parkes, L. M. The effect of black tea and caffeine on regional cerebral blood flow measured with arterial spin labeling. J Cereb Blood Flow Metab 2013; 33: 963–968.

118. Kim, W. S. H. et al. MRI Assessment of Cerebral Blood Flow in Nonhospitalized Adults Who Self-Isolated Due to COVID-19. Journal of Magnetic Resonance Imaging 2023; 58: 593–602.

119. Willie, C. K., Tzeng, Y.-C., Fisher, J. A. & Ainslie, P. N. Integrative regulation of human brain blood flow. J Physiol 2014; 592: 841–859.

120. Barton, C. W. & Wang, E. S. Correlation of end-tidal CO2 measurements to arterial PaCO2 in nonintubated patients. Ann Emerg Med 1994: 23: 560–563.

