## Supplementary Material for "Determinants of cerebral blood flow and arterial transit time in healthy older adults"

#### Methodology

##### 1: Participant inclusion criteria

Inclusion criteria were: 1) aged 60-85 yrs. 2) do <150 mins of moderate physical activity per week. 3) monolingual. 4) right-handed. 5) no current/historic diagnosis of cardiovascular, metabolic, respiratory, neurological, kidney, liver, or cancerous disease. 6) resting electrocardiogram (ECG) and blood pressure screened by clinician (i.e., no severe ECG abnormalities (e.g., ST depression, long QT, heart block, wide QRS) and systolic/diastolic blood pressure of <160/<90 mmHg, respectively). 7) Montreal Cognitive Assessment (MoCA) score ≥23. 8) not taking neurotransmitter-altering medication. 9) vaccinated against COVID-19. 9) deemed safe to enter MRI scanner by a qualified MRI-operator (i.e., absence/very small amount of ferrous metal in the body). 10) no language impairments. 11) no post-traumatic stress disorder (PTSD). 12) non-smoker of at least 5 yrs.

##### 2: Cardiorespiratory fitness test

***Treadmill test format example***

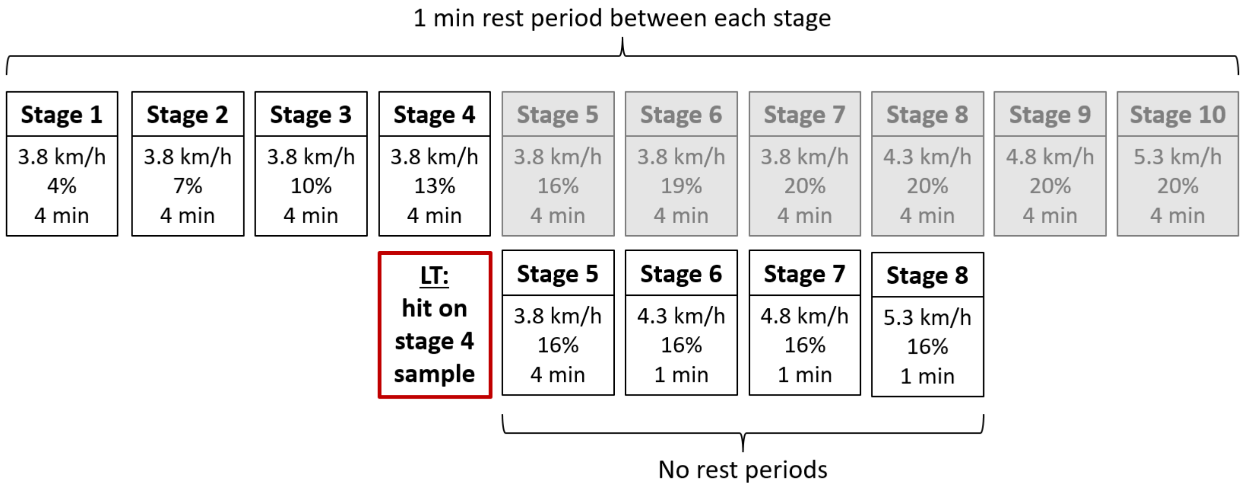

**Figure S1:** Incremental treadmill test format example where lactate threshold (LT) is hit after stage 4. Stages in grey are possible stages had lactate threshold not been hit after stage 4.

Oxygen consumption ($\dot{V}O_2$) and carbon dioxide production ($\dot{V}CO_2$) measured continuously. Lactate and RPE measured during each rest period and 1 min post-exercise. Heart rate recorded at the end of each stage. Participants began the next stage whilst waiting for lactate analysis from the previous rest period.

##### *Peak oxygen consumption prediction method*

Peak oxygen consumption ($\dot{V}O_{2peak}$) was predicted using the equation: x = (y-c)/m. Gradient (m) and intercept (c) were calculated from the line of best fit between three sub-maximal heart rate and $\dot{V}O_2$ data points from an individual's treadmill test, and y = age-predicted maximal heart rate ($HR_{age\text{-}pred}$; 220-age). This method assumes a largely linear relationship between heart rate and $\dot{V}O_2$ and that $HR_{age\text{-}pred}$ is relatively accurate (generally ±8–12 bpm).[1] This method was tested on 13 participants that completed high quality treadmill tests (peak values: $\%HR_{age\text{-}pred}$=103±5, respiratory exchange ratio (RER)= 1.14±0.03). The mean difference between actual ($\dot{V}O_{2peak}$) and predicted ($\dot{V}O_2$peak-pred) peak oxygen consumption was 8.6±2.5% (range=-11–12.3%). For n=8, the difference was <10%.

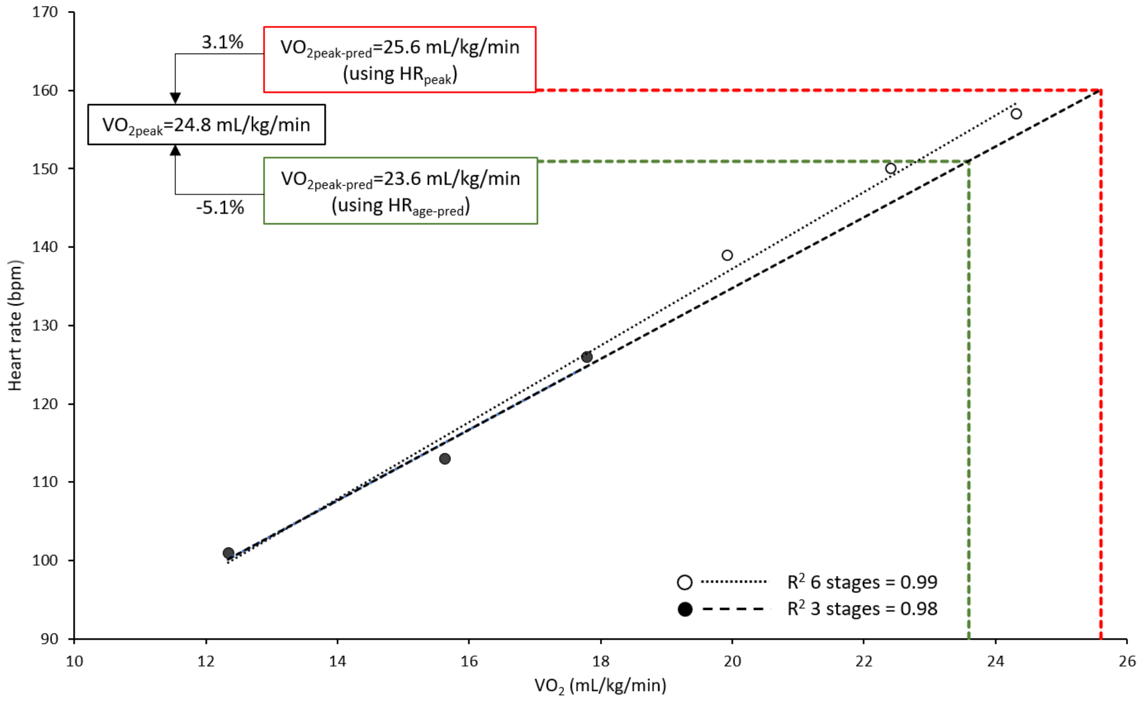

**Figure S2:** Example of peak oxygen consumption ($\dot{V}O_{2peak}$) prediction from sub-maximal heart rate and $\dot{V}O_2$ data (n=1).

The difference in the slope of the regression lines when using three and six sub-maximal stages is minimal. For this participant, predicted peak oxygen consumption ($\dot{V}O_{2peak\text{-}pred}$) was underestimated (-5.1%) when using $HR_{age\text{-}pred}$ (151 bpm; green line). For this participant, accuracy of this method was improved by 2% when using the peak heart rate ($HR_{peak}$) recorded in the treadmill test (160 bpm; red line). This method was validated in a sub-sample of participants who completed high quality treadmill test (n=13), mean change in predictive accuracy when using $HR_{age\text{-}pred}$ to $HR_{peak}$=2.1±3.4).

**3: MRI acquisition and analysis**

ASL data at each PLD (difference of tag and control averaged over repeats of each PLD) and grey matter masks in native space were visually inspected. Participants with abnormal ASL data were further investigated and n=6 data were excluded due to excessive motion. Grey matter masks were thresholded at 0.5 probability to ensure only voxels containing primarily grey matter were included in calculations of CBF and ATT. Areas within masks containing incorrect assignment to grey matter, primarily around the eyes and nasal cavity, were manually removed (n=7). Structural MRI data were aligned to the MNI brain using fsl_anat (https://fsl.fmrib.ox.ac.uk/fsl/fslwiki/fsl_anat). Registrations to MNI space were visually inspected. For participants with poor registration, the nonlinear registration was disabled and processing re-run. For participants where registration was deemed poor despite best efforts (due to brain atrophy with age), data were excluded from regional analysis requiring data in MNI space (n=1). Regional grey matter masks were made in MNI space and defined using the MNI structural atlas (temporal lobe only) or from the conjunction of the relevant regions from the Harvard atlas (in FSL). For n=9, MNI registration specifically of the inferior frontal lobe was poor (due to significant brain atrophy) and thus frontal lobe CBF and ATT maps were edited to exclude this region from analysis using fslroi.

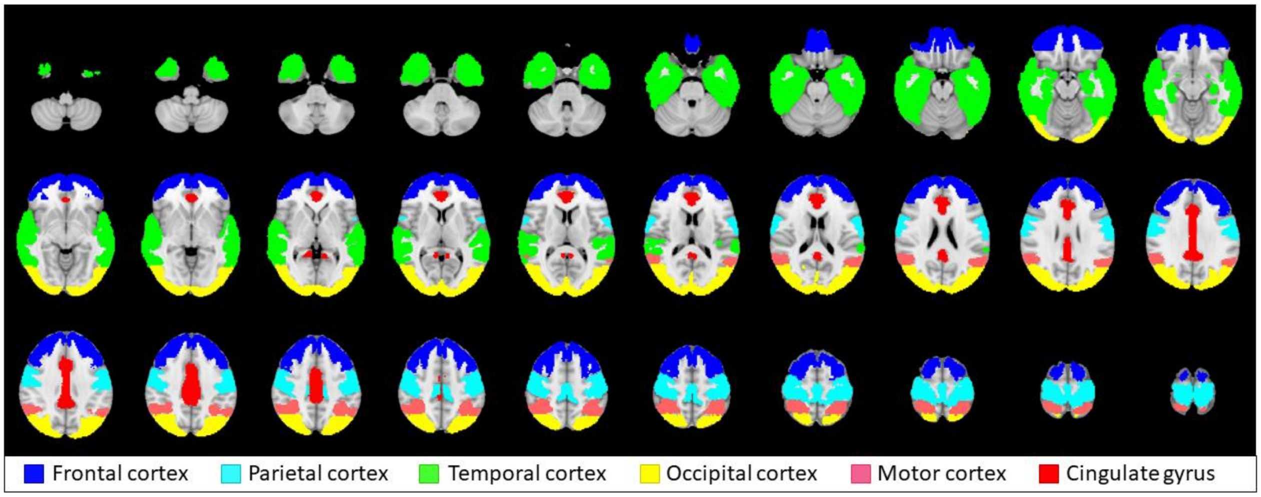

**Figure S3:** Grey matter masks used for region of interest analysis in MNI space.

### Results

##### 4: Global and regional data for CBF and ATT

**Table S1:** Global and regional values for grey matter CBF and ATT.

| | Total | Male | Female |
|---|---|---|---|
| **CBF (mL/100g/min)** | | | |
| Global | 63±12 | 62±12 | 65±12 |
| Frontal cortex | 78±17 | 78±21 | 79±18 |
| Parietal cortex | 85±19 | 85±23 | 86±19 |
| Temporal cortex | 56±12 | 55±15 | 58±11 |
| Occipital cortex | 78±18 | 75±21 | 81±18 |
| Motor cortex | 93±20 | 92±25 | 95±20 |
| Cingulate gyrus | 90±20 | 87±23 | 92±20 |
| **ATT (s)** | | | |
| Global | 1.42±0.17 | 1.43±0.17 | 1.40±0.17 |
| Frontal cortex | 1.37±0.14 | 1.39±0.26 | 1.35±0.14 |
| Parietal cortex | 1.50±0.16 | 1.54±0.30 | 1.46±0.15 |
| Temporal cortex | 1.26±0.13 | 1.27±0.24 | 1.26±0.13 |
| Occipital cortex | 1.59±0.19 | 1.63±0.32 | 1.55±0.19 |
| Motor cortex | 1.43±0.14 | 1.45±0.27 | 1.40±0.14 |
| Cingulate gyrus | 1.25±0.18 | 1.25±0.26 | 1.25±0.19 |

*Values represent means ± standard deviation. Global (n=78) and regional (n=77) values are from native and MNI space, respectively. CBF; cerebral blood flow, ATT; arterial transit time.*

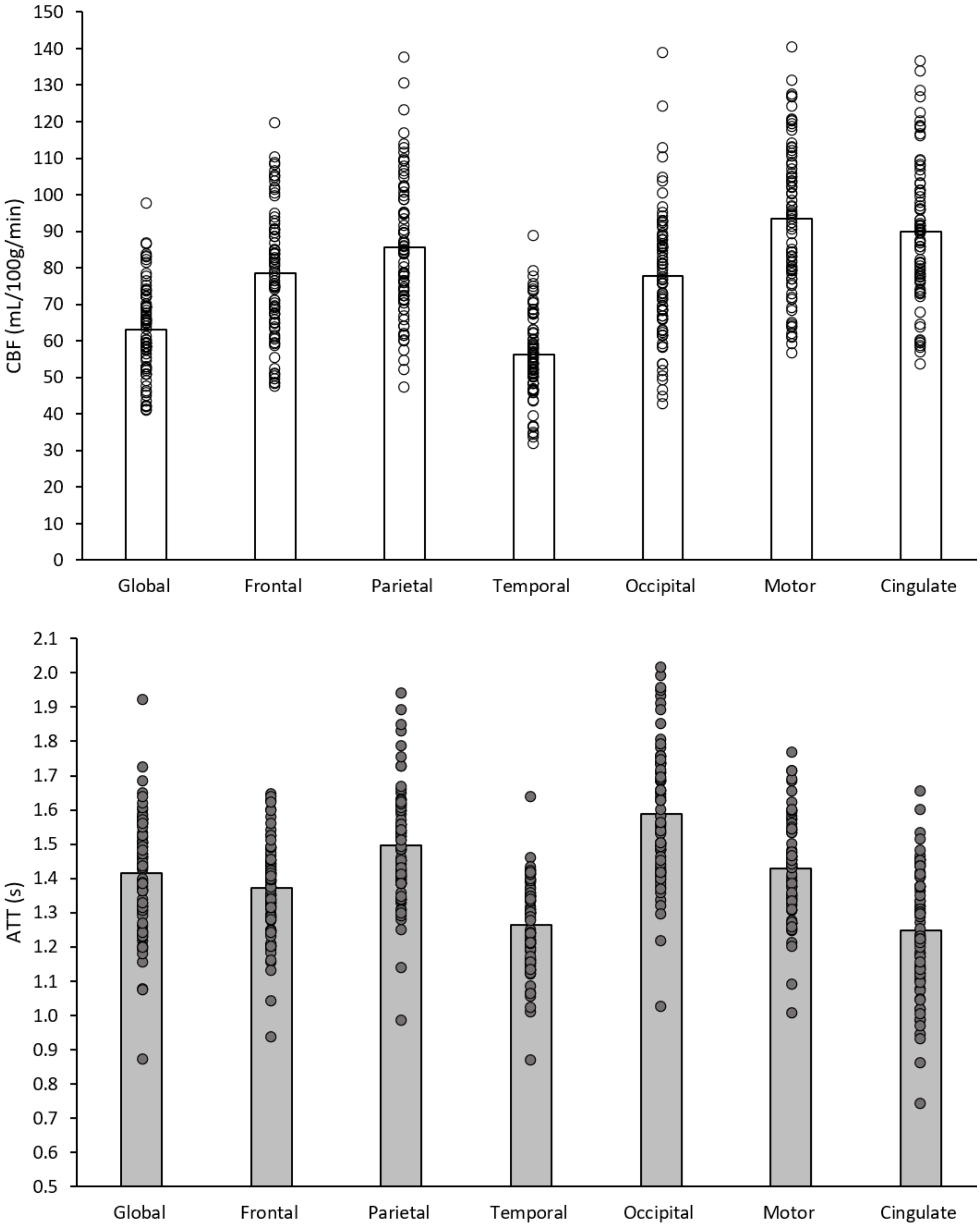

**Figure S4:** Means and individual data for global and regional cerebral blood flow (CBF; top) and arterial transit time (ATT; bottom) in healthy older adults. Global (n=78) and regional (n=77) analyses were performed in native and MNI space, respectively.

**Table S2:** Associations between age, BMI, and cardiorespiratory fitness with regional CBF.

| | Age | | BMI | | $\dot{V}O_{2peak}$ | |
|---|---|---|---|---|---|---|
| **CBF** | β | *P* | β | *P* | β | *P* |
| Frontal | -0.07 | 0.613 | -0.34 | **0.008** | -0.16 | 0.346 |
| Parietal | -0.05 | 0.722 | -0.41 | **0.002** | -0.13 | 0.446 |
| Temporal | -0.12 | 0.361 | -0.44 | **<0.001** | -0.20 | 0.213 |
| Occipital | -0.06 | 0.639 | -0.43 | **<0.001** | -0.08 | 0.597 |
| Motor | -0.03 | 0.805 | -0.35 | **0.007** | -0.19 | 0.257 |
| Cingulate | 0.00 | 0.987 | -0.39 | **0.002** | -0.11 | 0.515 |

*Separate multiple linear regressions were performed for each region (n=77), independent variables: age, sex, BMI, and $\dot{V}O_{2peak}$. Bold indicates significant P values survived adjustment for multiple comparisons.* β*; standardised beta coefficient, CBF; cerebral blood flow, BMI; body mass index, $\dot{V}O_{2peak}$; peak oxygen consumption.*

**5: Associations between cognitive function and CBF or ATT**

***Global analysis***

**Table S3:** Associations between global CBF or ATT with cognitive function.

| | **gCBF (mL/100g/min)** | | **gATT (s)** | |
|---|---|---|---|---|
| n=76 | β | *P* | β | *P* |
| Age (years) | 0.113 | 0.400 | 0.325 | 0.015 |
| Sex | 0.096 | 0.435 | -0.107 | 0.372 |
| Education | -0.079 | 0.522 | 0.075 | 0.532 |
| *Processing speed* | | | | |
| Response time | -0.049 | 0.696 | -0.001 | 0.995 |
| Accuracy | 0.247 | 0.056 | -0.082 | 0.514 |
| *Working memory* | | | | |
| 2-back *d* prime | 0.170 | 0.190 | 0.016 | 0.900 |
| *Attention (response time)* | | | | |
| Alerting | -0.122 | 0.344 | 0.041 | 0.746 |
| Orienting | 0.073 | 0.580 | -0.017 | 0.896 |
| Executive control | 0.111 | 0.405 | -0.034 | 0.795 |

*Results from two multiple linear regression analyses. β; standardised beta coefficient, gCBF; global cerebral blood flow, gATT; global arterial transit time. Sex (1; male, 2; female). Education (1; compulsory, 2; further, 3; undergraduate, 4; post-graduate).*

***Regional analysis***

To analyse regional differences, multiple linear regressions were performed with regional CBF or ATT as dependent variables, and age, sex, education (1; compulsory, 2; further, 3; undergraduate, 4; post-graduate) and scores for processing speed (accuracy and response time), working memory (*d* prime), and the three attentional domains (alerting, orienting, and executive control response times) as independent variables. MNI registration was poor for n=1 and cognitive data was missing for n=2, leaving n=75 for regional analysis.

All associations between CBF or ATT of any region and any cognitive measures were non-significant. Only processing speed accuracy showed any indication of a trend with non-significant positive associations with CBF in all regions ($\beta$=0.23–0.25, *P*=0.052–0.080)

### References

1. Achten, J. & Jeukendrup, A. E. Heart Rate Monitoring: Applications and Limitations. *Sports Medicine* 2003; 33 517–538
